# LIFU-Responsive Nanocomplexes Deliver PDGF-BB mRNA for Plaques Stabilization via Neovascularization Modulation

**DOI:** 10.64898/2026.07.23.740436

**Authors:** Qirong Xie, Xinxin Li, Liming Deng, Yue Ran, Xun Yang, Fan Liu, Yu Chen, Jianhao Luo, Shujuan Su, Daofa Zhang, Dan Deng, Qunxia Zhang, Jianli Ren, Zhigang Wang, Haitao Ran, Rongzhong Huang, Chunyan Ma, Yang Sun

## Abstract

**Background:** Pathological intraplaque neovascularization, vascular leakage, and fibrous cap thinning contribute to vulnerable atherosclerotic plaque rupture. Platelet- derived growth factor-BB (PDGF-BB) has been shown to promote pericyte recruitment, thereby stabilizing the microvascular structure, and to induce phenotypic modulation of vascular smooth muscle cells (VSMCs), which enhances fibrous cap thickness and reinforces plaque stability. Nevertheless, systemic protein delivery is limited by rapid clearance and potential off-target effects.

**Methods:** We developed PDGF-BB mRNA-loaded lipid nanoparticle–poly(lactic-co- glycolic acid) nanobubble complexes (LNPmRNA@PLGA) and used low-intensity focused ultrasound (LIFU) to enhance plaque-targeted delivery. Cellular uptake, PDGF- BB expression, vascular mural-cell responses, plaque histology, hemodynamics, and proteomic changes were evaluated in vitro and in ApoE^−/−^Fbn1^C1041G+/−^ mice.

**Results:** LIFU enhanced nanocomplex uptake and PDGF-BB expression, promoted vascular smooth muscle cell proliferation, migration, and phenotypic switching, and increased pericyte coverage. In vivo, LIFU plus LNPmRNA@PLGA reduced the plaque vulnerability index by 78.2% and the neovascularization area by 67.3% compared with controls, while increasing collagen deposition and improving carotid hemodynamics.

**Conclusions:** LIFU-responsive delivery of PDGF-BB mRNA stabilized vulnerable plaques by promoting neovessel maturation and strengthening the fibrous cap. This strategy provides a spatially controlled framework for therapeutic remodeling of high-risk atherosclerotic plaques.

**Research Perspective:** *What New Question Does This Study Raise?:*

- Can spatially controlled PDGF-BB mRNA delivery simultaneously mature intraplaque neovessels and reinforce the fibrous cap without the systemic effects associated with recombinant PDGF-BB?

*What Question Should Be Addressed Next?:*

- Future studies should define the therapeutic window, durability, and long-term safety of LIFU-triggered PDGF-BB mRNA delivery in large-animal models that more closely reproduce human plaque rupture.

## Introduction

Atherosclerotic plaque rupture underlies the majority of acute myocardial infarctions and ischemic strokes, yet current pharmacological treatments remain of limited efficacy.^1,2^ Growing evidence suggests that plaque destabilization and rupture are driven by immature and leaky neovessels, which compromise vascular integrity. Significantly, leakage from these neovessels contributes to local hypoxia and inflammation, facilitating the infiltration of erythrocytes, inflammatory mediators, and lipids into the plaque, which promotes matrix degradation, fibrous cap thinning, and increased plaque vulnerability.^1,3^ Given the presence of immature and dysfunctional neovessels in vulnerable plaque, normalization of intraplaque neovessels may represent a promising therapeutic target for advanced atherosclerosis.

Among the cells involved in atherosclerotic neovascularization, mural cells—including pericytes and vascular smooth muscle cells (VSMCs)—serve as critical regulators of neovessel maturation and stability.^4^ Platelet-derived growth factor-BB (PDGF-BB) stabilizes intraplaque neovessels by recruiting mural cells,^5–9^ while simultaneously promoting the phenotypic switching of VSMCs from a contractile to a synthetic state and enhancing collagen deposition within the fibrous cap^10,11^, thereby contributing to plaque stabilization.^12,13^ Despite its therapeutic potential, preclinical studies report suboptimal outcomes, largely due to uncontrolled PDGF-BB release, as excessive PDGF-BB may lead to plaque overgrowth and exacerbate hypoxia.^14^ In addition, although recombinant PDGF-BB has been clinically applied for local or topical use, its systemic application in atherosclerosis is hindered by rapid clearance and potential off-target effects such as fibrosis.^15,16^

mRNA-based therapies may offer an attractive alternative, as they enable rapid and efficient in vivo production of therapeutic proteins.^17–19^ Yet, to fully exploit this advantage, mRNA requires a delivery system capable of ensuring targeted and imaging-guided transport to protect the payload and facilitate cellular uptake. 7C1, an oligomer-lipid conjugate, exhibits high affinity for endothelial cells through optimized pKa control, minimal ApoE adsorption, hydrophobic C15 chains, and caveolae-mediated transcytosis.^20,21^ As reported, it efficiently mediates mRNA transfection in endothelial cells, including those in atherosclerotic plaques.^22^

Therefore, in our study, PDGF-BB mRNA was first encapsulated within 7C1 cationic liposomes and electrostatically attached to PLGA nanobubbles to form LNPmRNA@PLGA nanocomplexes. Our previous studies have shown that poly(lactic-co-glycolic acid) (PLGA) nanobubbles (NBs) act as ultrasound-responsive carriers that can be disrupted under low-intensity focused ultrasound (LIFU) exposure to enhance cellular uptake of therapeutic agents.^23–26^ Accordingly, LIFU exposure was used to facilitate delivery of the encapsulated mRNA into endothelial cells. This increased local PDGF-BB expression, modulated pericyte recruitment and VSMC behavior, promoted neovessel maturation, enhanced fibrous cap integrity, and ultimately improved plaque stability. Overall, these results demonstrate that our strategy has the potential to stabilize vulnerable plaques by remodeling the intraplaque neovascular network.

## Methods

Preparation of LNPmRNA@PLGA: LNPmRNA complexes were prepared via microfluidic mixing of 7C1 lipid (1.5 mg, SunLipo NanoTech) and C14-PEG2000 (0.5 mg, Ponsure) dissolved in anhydrous ethanol, with PDGF-BB mRNA (The mRNA sequences are provided in Table S1) dissolved in sterile sodium citrate buffer (50 mM, pH 4.0). PLGA nanobubbles (NBs) were synthesized using a double emulsion technique: PLGA polymer (5 mg, MCE) was dissolved in dichloromethane (100 µL) under vortex mixing. Perfluoropentane (PFP) (20 µL, Aladdin) was then emulsified with the PLGA solution through acoustic oscillation (30 s) to form a primary emulsion. This emulsion was stabilized by adding 4% polyvinyl alcohol (500 µL) and homogenized at 12,000 rpm for 5 min, followed by solvent evaporation using 2% isopropanol (1 mL). Final LNPmRNA@PLGA nanocomplexes were assembled at a mass ratio of 16:1 (7C1 cationic lipid to PLGA) in phosphate-buffered saline (PBS, pH 5.0), incubated at room temperature for 30 minutes, and subsequently subjected to overnight dialysis against PBS (pH 7.4).

Optimal binding rate between LNPmRNA and PLGA gas core: Fluorescent lipid phase was prepared by adding 1.5 μL Cy5.5 (or DiI for microscopy) to ethanol to a final concentration of 10 mg/mL, and used to formulate LNPmRNA via microfluidics. PLGA cores were labeled with 5 μL FITC (10 mg/mL in PBS), incubated at 25 °C for 30 min in the dark, and centrifuged (15,000×g, 20 min) to remove unbound dye. Labeled LNPmRNA and PLGA cores were mixed at mass ratios 0.5:1-32:1 for electrostatic self- assembly to form LNPmRNA@PLGA nanocomplexes. The optimal ratio was determined by quantifying dual-fluorescence via nano-flow cytometry, and colocalization was confirmed by confocal laser scanning microscopy (Zeiss LSM 980). All experiments were repeated in triplicate.

Characterization of LNPmRNA@PLGA: The morphological features of LNPmRNA@PLGA nanocomplexes were analyzed by transmission electron microscopy (TEM, Hitachi HT-7800) operating at 120 kV. Particle size distribution was quantified using dynamic light scattering (DLS) with a Malvern Zetasizer Nano ZS (Malvern Panalytical), while surface charge characteristics were determined through phase analysis light scattering (PALS) measurements on the same instrument. To assess the stability of LNPmRNA@PLGA, the nanocomplexes were incubated in PBS or medium containing 10% FBS at 4°C or 37°C, and changes in particle size were monitored over 7 or 2 days. To evaluate mRNA encapsulation efficiency, both bare LNPmRNA and core- shell LNPmRNA@PLGA formulations were treated with 2% (v/v) Triton X lysis buffer (Sigma-Aldrich) for 15 min at 37°C. The released mRNA was electrophoresed on a 1% (w/v) agarose gel (120 V, 30 min) containing GelRed nucleic acid stain (Biotium).

The Effect of Temperature on the In Vitro Imaging Performance of LNPmRNA@PLGA: Five temperature groups were set at 25°C, 30°C, 35°C, 37°C, and 40°C. The LNPmRNA@PLGA nanocomplexes were injected into the gel models, and the respective gels were heated while monitoring the temperature with an infrared camera. Once the internal gel temperature reached the target temperature, the imaging performance in B-mode and contrast mode was immediately observed using the Vevo® LAZR system.

In Vitro Ultrasound Imaging of LNPmRNA@PLGA: Weigh 10.5 g of agarose powder and dissolve it in 300 mL of distilled water. Stir the mixture until fully dissolved, then heat it in a microwave at high power for 3 minutes until the agarose solution becomes clear and transparent. Pour the hot agarose solution into a 1.25 mL pipette tip box to prepare the in vitro gel model. LNPmRNA@PLGA nanocomplexes were uniformly dispersed within the gel matrix. Phase transition was induced by localized heating to 37°C (±0.5°C) using a thermostatic water bath (Julabo, 5 min equilibration). Ultrasound-responsive behavior was quantified using a high-resolution imaging system (Vevo® LAZR, Visual Sonics) equipped with an LZ250D transducer (21 MHz center frequency). In Vivo Ultrasound Imaging of LNPmRNA@PLGA: ApoE^−/−^Fbn1^C1041G+/−^ mice were injected with LNPmRNA@PLGA nanocomplexes (1mg/kg) via the tail vein. Realtime ultrasound imaging of the right carotid artery was then performed using an ultrasound system (VINNO ultimus 9E) to monitor the contrastenhanced effect of the nanocomplexes.

Biological Safety of LNPmRNA@PLGA: Male C57BL/6 mice (6 w), obtained from Chongqing Medical University, were randomized into four groups (n = 3 per group): saline (control) and mRNA concentration at doses of 0.25, 0.5, and 1 mg/kg, administered via intravenous injection. At 24 h post-injection, mice were euthanized, and blood and major organs were collected for biochemical and histological analyses, respectively.

In Vivo Biodistribution of LNPmRNA@PLGA: Male C57BL/6 mice (approximately 20 g) were randomized into three groups (n = 3 per group): control (PLGA NBs), Cy7.5- labeled PLGA NBs, and Cy7.5-labeled LNPmRNA@PLGA nanocomplexes. All formulations were administered via tail vein injection at a dose of 1 mg/kg. At 2, 4, 6, and 8 h post-injection, major organs (heart, lung, liver, spleen, and kidney) were harvested and imaged using an in vivo fluorescence imaging system (Kodak, USA).

Endothelial Tropism Evaluation of LNPmRNA@PLGA: To evaluate the endothelial tropism of LNPmRNA@PLGA, six 20 weeks ApoE^−/−^Fbn1^C1041G+/−^ mice were randomized into two groups (n=3 each): Cy5.5-LNPmRNA@PLGA and LIFU+Cy5.5- LNPmRNA@PLGA. Both groups received the nanocomplexes via tail vein injection (1 mg/kg), and the LIFU group received immediate LIFU irradiation (1 W/cm², 4 min) postinjection. After sacrifice, hearts and aortas were harvested. Aortas were imaged via smallanimal fluorescence imaging (Kodak, USA), and aortic root sections were stained for immunofluorescence to assess nanoparticle accumulation in plaques. Cell Culture: HUVECs (Human umbilical vein endothelial cells)were cultured in Dulbecco’s modified eagle medium supplemented with 10% fetal bovine serum (Pricella, China), C3H/10T1/2 cells and VSMCs (Mouse derived vascular smooth muscle cells) were cultured in RPMI-1640 medium supplemented with 20% fetal bovine serum (GIBCO, CA), Cells were incubated at 37°C and 5% carbon dioxide.

Cell cytotoxicity assessment: HUVECs (5×10³/well) were seeded in 96-well plates and treated with LNPmRNA@PLGA at various concentrations (mRNA: 1, 2, and 4 μg/mL). After 4 h of pre-culture at 37°C in 5% CO₂, nanocomplexes were added and incubated for 24 h. The medium was replaced with 100 μL of fresh medium containing 10% CCK-8, followed by a 4 h incubation. Absorbance at 450 nm was measured using a microplate reader (BioTek).

Effect of LIFU on the Viability of HUVECs: HUVECs (5×10³ cells/well in 96-well plates) were divided into control and LIFU-treated groups (1, 2, 4 W/cm²) with irradiation durations of 0-8 min (n=5). Post-irradiation cultures were maintained for 24 h, followed by CCK-8 assay: 100 μL medium with 10% reagent was added per well, incubated 4 h, and absorbance measured at 450 nm (BioTek microplate reader).

Cellular uptake analysis under LIFU stimulation: HUVECs (1×10⁶ cells/dish) were plated on glass-bottom dishes and allowed to adhere for 12 h. Cells were treated with DiI-labeled LNPmRNA@PLGA (4 μg/mL mRNA equivalence) for 6 h, with parallel LIFU irradiation (parameters as established) applied to the experimental group. Post-treatment, cells were fixed with 4% PFA and nuclei counterstained with DAPI. Subsequently, cellular internalization efficiency was evaluated by flow cytometry, and the intracellular distribution of the nanoparticles was visualized using confocal laser scanning microscopy.

PDGF-BB mRNA and protein expression analysis: HUVECs in 6-well plates were treated with nanoparticle formulations (4 h), followed by 24 h incubation in fresh medium. Total RNA was isolated using TRIzol (Beyotime), reverse-transcribed into cDNA, and analyzed via qPCR (CFX96, Bio-Rad) with PDGF-BB-specific primers (Table S2). ΔΔCt method normalized expression to GAPDH. HUVECs treated with nanoparticle groups (protocol as above) were lysed after 24 h. Total protein (20 μg/lane by BCA assay) was resolved via SDS-PAGE, transferred to PVDF membranes, and probed with anti-PDGF- BB (Abcam, 1:1000) and GAPDH (loading control). Signals were developed using ECL (Bio-Rad).

PDGF-BB expression in Plaques: To evaluate PDGF-BB expression in atherosclerotic plaques, fifteen 8-week-old ApoE^−/−^Fbn1^C1041G+/−^ mice were randomly divided into five groups (n = 3): Control, LNPmRNA, LIFU+LNPmRNA, LNPmRNA@PLGA, and LIFU+LNPmRNA@PLGA. All treatments were administered via tail vein injection over 4 weeks. Subsequently, mice were euthanized and aortic roots were harvested for immunofluorescence staining of PDGF-BB. The relative PDGF-BB content was quantified as the positive staining area normalized to total plaque area.

The proliferation of C3H/10T1/2 cells and VSMCs: Conditioned medium (CM) from nanoparticle-treated HUVECs (48 h culture, 0.22 μm filtered) was applied to C3H/10T1/2 fibroblasts and VSMCs (5×10³ cells/well, n=5). Cells were exposed to PDGF-BB-enriched CM for 24 h, followed by CCK-8 viability assessment (protocol as above).

The migration ability of C3H/10T1/2 cells and VSMCs: C3H/10T1/2 fibroblasts and VSMCs (3×10⁴ cells/insert) were seeded in Transwell chambers (8 μm pore, Corning) with PDGF-BB-enriched medium (500 μL) in the lower chamber. After 48 h incubation, cells were fixed with 4% PFA, stained with 0.1% crystal violet (Sigma), and quantified by counting five random fields/membrane (Nikon Eclipse Ti, 20× objective).

Flow cytometry experiment: VSMCs were seeded in six-well plates by group. After adhesion, cells were treated with PDGF-BB-containing medium for 24 hours, then digested into single-cell suspensions (1 × 10⁶ cells/mL). Contraction- and synthesis- phenotype populations were analyzed by flow cytometry following centrifugation and incubation with anti-α-SMA and anti-OPN-1 antibodies.

Confocal microscopy experiment: VSMCs were seeded in confocal dishes by group. Post-adhesion, cells were treated with PDGF-BB for 24 h, fixed with 4% paraformaldehyde (20 min), permeabilized with 0.1% Triton X-100 (15 min), and incubated with anti-α-SMA and anti-OPN-1 antibodies (45-60 min). After PBS washes, protein staining was imaged by confocal microscopy and statistically analyzed.

Construction of the vulnerable plaque mouse model: Vulnerable plaque mice (ApoE^−/−^Fbn1^C1041G+/−^) were generated by crossing Fbn1^C1041G+/−^ and ApoE^−/−^ strains (Shanghai Southern Model Organisms) and fed a high-fat diet for 12 weeks from 8 weeks of age. After the model mice were successfully constructed, the genotypes of the model mice were identified. Genomic DNA from mouse tail samples was extracted (Beyotime kit), amplified with Fbn1^C1041G+/−^ and ApoE^−/−^ primers (Table S2), and analyzed by gel electrophoresis and sequencing.

Histological analysis of mouse arteries: Aortic roots from treated mice were sectioned for H&E, Oil Red O(ORO), and Sirius Red staining. Immunohistochemistry (TER-119, Biosciences) and immunofluorescence (CD31/NG2/α-SMA/OPN-1/CD68/HIF-1α, Servicebio) were performed on aortic sections.

ELISA analysis of plaque tissue and serum: Aortic plaque tissues were rinsed with ice- cold PBS, weighed, homogenized in PBS containing protease inhibitors, and centrifuged at 4°C. The resulting supernatants were collected, and total protein concentrations were determined for normalization. CD68 and HIF-1α levels in plaque- tissue lysates were measured using enzyme-linked immunosorbent assay (ELISA) kits according to the manufacturers’ instructions. Serum was isolated from whole blood by centrifugation, and circulating TNF-α and IL-6 concentrations were measured using the corresponding ELISA kits. Absorbance was recorded with a microplate reader, and concentrations were calculated from standard curves.

Carotid ultrasound and hemodynamic assessment: Mice were anesthetized and placed supine on a temperature-controlled platform. After removal of cervical hair and application of prewarmed coupling gel, the right common carotid artery (RCCA) and right internal carotid artery (RICA) were visualized using high-frequency B-mode and color Doppler ultrasound (VINNO 6 LAB). Pulsed-wave Doppler spectra were acquired with the sample volume positioned in the center of the vessel lumen and the insonation angle maintained at ≤60°. Peak systolic velocity (PSV) and resistive index (RI) were measured over at least three consecutive cardiac cycles. Measurements were repeated three times for each vessel, and the mean value was used for statistical analysis.

Data-independent acquisition proteomics and bioinformatic analysis: Aortic tissues from the control and LIFU + LNPmRNA@PLGA groups were collected for proteomic analysis. Total proteins were extracted, quantified, enzymatically digested into peptides, and analyzed by liquid chromatography–tandem mass spectrometry using a data-independent acquisition (DIA) workflow. Differentially abundant proteins were defined using *P* < 0.05 and |log2(fold change)| ≥ 1. Principal component analysis (PCA) was used to assess separation between groups. Gene Ontology (GO) and Kyoto Encyclopedia of Genes and Genomes (KEGG) enrichment analyses were performed to identify overrepresented biological processes and pathways, and Gene Set Enrichment Analysis (GSEA) was used to evaluate coordinated pathway-level changes.

Statistical analysis: All data are presented as mean ± SD. Statistical significance (*P* <0.05, *P* < 0.01, *P* < 0.001, *P* < 0.0001) was determined by Unpaired t test for (two groups) or one-way ANOVA with Tukey’s test (multi-group), validated by ≥3 independent experiments. Analyses used SPSS 25/GraphPad Prism 9.5.1.

## Results

### Characterization of LNPmRNA@PLGA

The preparation of LNPmRNA@PLGA NPs is illustrated in Scheme 1. LNPmRNA and PLGA NBs were synthesized using microfluidic and double emulsion methods, respectively. LNPmRNA@PLGA nanocomplexes were then fabricated by the hetero- assembly of negatively charged LNPmRNA (-13.3 ± 0.85 mV) and positively charged PLGA NBs (+5.6 ± 0.45 mV) at pH 5.0. Nanoflow cytometry and fluorescence microscopy were used to determine the optimal LNPmRNA: PLGA mass ratio, which was found to be 16:1 and adopted for subsequent experiments (Fig. 1A and Fig. S1A, B). As shown in Fig. 1B, the average diameter of LNPmRNA@PLGA (484 ±25.58 nm) was slightly incresed compared with PLGA NBs alone (316 ± 6.27 nm), with comparable PDI values. The electrostatic interaction between negatively charged LNPmRNA and positively charged PLGA NBs partially neutralized the surface charge, resulting in a near-neutral zeta potential of − (0.86 ± 0.20 mV) for LNPmRNA@PLGA. Transmission electron microscopy (TEM) (Fig. 1C) images revealed that both LNPmRNA and PLGA NBs were spherical and uniformly dispersed. In LNPmRNA@PLGA nanocomplexes, LNPmRNA was attached to the surface of PLGA NBs, confirming the successful assembly of LNPmRNA@PLGA nanocomplexes.

**Figure 1.**
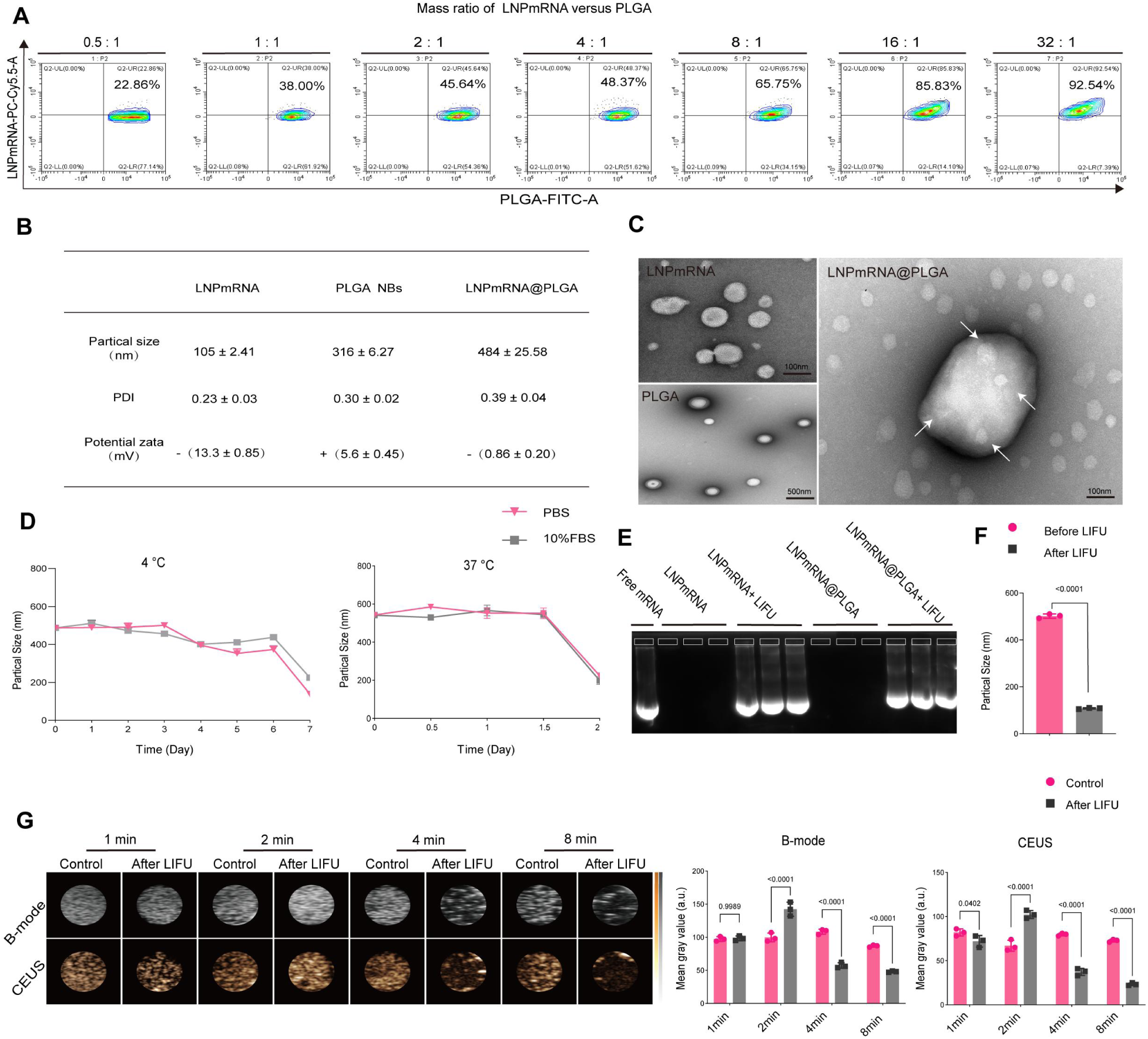
Preparation and Characterization of LNPmRNA@PLGA Nanocomplexes. (A) Optimal mass ratio between LNPmRNA and PLGA NBs determined by nanoflow cytometry. (B) Particle size, polydispersity index (PDI), and zeta potential of LNPmRNA, PLGA NBs, and LNPmRNA@PLGA. (C) Transmission electron microscopy images of LNPmRNA, PLGA NBs, and LNPmRNA@PLGA. (D) Particle size changes of LNPmRNA@PLGA during incubation in PBS or 10% FBS at 4°C and 37°C. (E) Gel electrophoresis analysis of mRNA protection and LIFU-triggered mRNA release from LNPmRNA and LNPmRNA@PLGA. (F) Mean particle size of LNPmRNA@PLGA before and after LIFU irradiation. (G) Ultrasound imaging of LNPmRNA@PLGA after LIFU irradiation for different durations. Quantification of ultrasound signal intensity in B-mode and contrast-enhanced ultrasound (CEUS) mode. Data are presented as mean ± SD. An unpaired t test was used for (F) and (G).

Subsequently, the in vitro stability of LNPmRNA@PLGA was evaluated. LNPmRNA@PLGA nanocomplexes were incubated in PBS or 10% fetal bovine serum (FBS) at either 4°C or 37°C, and changes in particle size were monitored over 7 days. As shown in Fig. 1D, LNPmRNA@PLGA remained stable for up to 6 days at 4°C and for 1.5 days at 37°C, demonstrating adequate suitability for in vivo applications.

The mRNA-protective capability of LNP was evaluated by gel electrophoresis. As shown in Fig. 1E, LNP effectively prevented mRNA degradation while allowing mRNA release after LIFU exposure. LNPmRNA@PLGA exhibited comparable mRNA protection to LNP alone, confirming its suitability for mRNA delivery. Upon LIFU irradiation, the particle size of LNPmRNA@PLGA decreased from (501 ± 9.07 nm) to (108 ± 2.64 nm), further demonstrating that LIFU effectively triggered nanocomplex dissociation and facilitated mRNA release (Fig. 1F).

### Parameter optimization and biosafety assessment of LIFU combined with LNPmRNA@PLGA

To optimize LIFU parameters while ensuring biosafety, the viability of human umbilical vein endothelial cell (HUVEC) was evaluated using a CCK-8 assay. As shown in Fig. S1D, cell viability decreased with increasing acoustic intensity and irradiation duration. Specifically, HUVEC viability remained approximately 89% after 8 min of LIFU irradiation at 1 W/cm², indicating acceptable biosafety under this condition. In contrast, viability progressively declined at higher acoustic intensities, reaching 68% and 52% after 8 min exposure of LIFU at 2 and 4 W/cm², respectively. Based on these findings, ultrasound imaging was subsequently performed to optimize irradiation duration. As shown in Fig. 1G, ultrasound contrast signal rapidly increased within the first 2 min of irradiation and gradually declined thereafter. This transient enhancement was likely attributable to the LIFU induced liquid-to-gas transition of encapsulated PFP, which generated nanobubbles and enhanced acoustic impedance mismatch. Prolonged irradiation led to progressive nanobubble collapse and signal attenuation. Accordingly, 1 W/cm² for 4 min was selected as the optimal LIFU condition, providing a balance between biosafety (HUVEC viability >85%) and sustained contrast performance (nanobubble stability >75%).

### Ultrasound Imaging and Targeting Capability of LNPmRNA@PLGA

Perfluoropentane (PFP), a thermally responsive material, plays a critical role in LNPmRNA@PLGA-mediated ultrasound imaging through its phase-change capability. To determine the optimal phase-change condition, the ultrasound imaging performance of LNPmRNA@PLGA was evaluated at different temperatures. As shown in Fig. 2A, the strongest contrast enhancement was observed at 37°C in both B-mode and contrast enhanced ultrasound modes, indicating efficient phase transition under physiological conditions. At 37°C, PFP underwent liquid-to-gas phase transition and generated nanobubbles, thereby increasing acoustic impedance mismatch and acoustic backscatter. The temporal stability of ultrasound imaging was then evaluated. After injection into 37°C-preheated gel phantoms (Fig. 2B), LNPmRNA@PLGA nanocomplexes maintained imaging intensity for up to 15 min, followed by gradual signal attenuation (Fig. 2C). Consistent with the in vitro findings, in vivo dynamic imaging showed sustained contrast enhancement in murine carotid arteries after LNPmRNA@PLGA administration (Movie S1). Together, these results indicate that the LNPmRNA@PLGA provides stable spatiotemporal ultrasound contrast, supporting its use for ultrasound-guided in vivo visualization.

**Figure 2.**
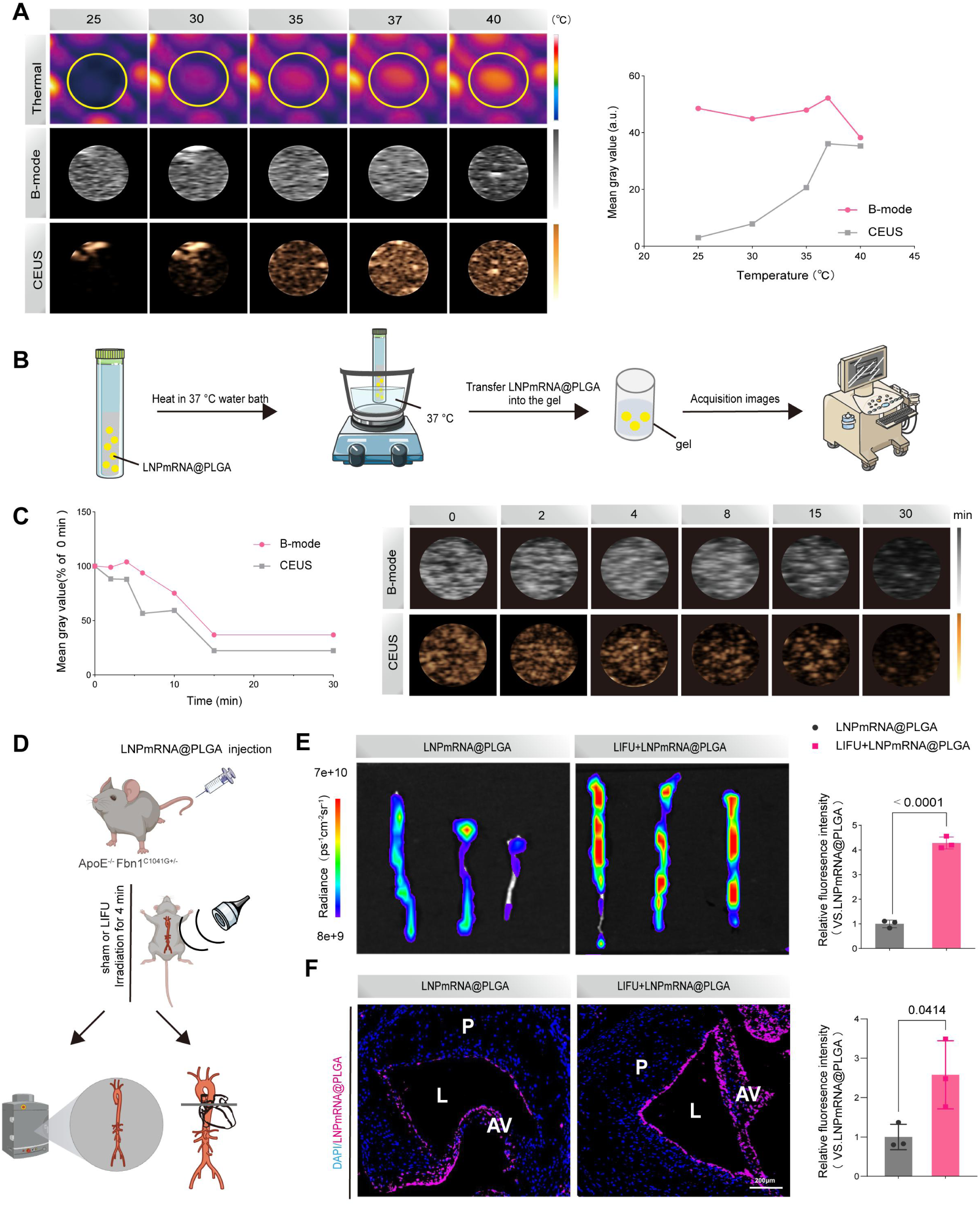
Ultrasound imaging performance and plaque-targeting capability of LNPmRNA@PLGA. (A) Temperature-dependent phase-transition imaging of LNPmRNA@PLGA. (B) Schematic illustration of the in vitro ultrasound imaging procedure for LNPmRNA@PLGA. (C) Time-dependent ultrasound imaging and echo signal quantification of LNPmRNA@PLGA. (D) Schematic illustration of LNPmRNA@PLGA administration and LIFU-guided targeting in ApoE^−/−^Fbn1^C1041G+/−^ mice. (E) Ex vivo fluorescence imaging of the aorta after treatment with LNPmRNA@PLGA alone or LIFU + LNPmRNA@PLGA. (F) Immunofluorescence images of aortic root plaques showing LNPmRNA@PLGA accumulation. P, plaque; L, lumen; AV, aortic valve. An unpaired t test was used for (E) and (F).

The LIFU-enhanced plaque-targeting capability of LNPmRNA@PLGA nanocomplexes was evaluated in ApoE^−/−^Fbn1^C1041G+/−^ mice. As illustrated in Fig. 2D, mice received intravenous injections with LNPmRNA@PLGA and then randomized into LIFU-treated or untreated group (1 W/cm², 4 min). After treatment, the aortas were harvested for ex vivo fluorescence imaging, followed by immunofluorescence analysis of aortic root plaques to confirm nanocomplex accumulation. As shown in Fig. 2E and F, detectable fluorescence signals were observed in the arteries of mice receiving LNPmRNA@PLGA, and immunofluorescence analysis further confirmed nanocomplex localization within atherosclerotic plaques. Notably, LIFU irradiation significantly increased nanocomplex accumulation, as evidenced by stronger fluorescence signals in both ex vivo aortic imaging and aortic root sections. These findings indicate that LIFU enables spatiotemporally controlled delivery of nanocomplexes to atherosclerotic plaques.

To further assess the systemic biodistribution and clearance profile of LNPmRNA@PLGA, fluorescence imaging was performed (Fig. S2). As expected, unlabeled PLGA NBs exhibited negligible fluorescence signals, confirming minimal background interference. Fluorescence signals from both Cy7.5-labeled PLGA NBs and LNPmRNA@PLGA nanocomplexes were predominantly detected in the liver and kidneys. Compared with Cy7.5-labeled PLGA NBs, LNPmRNA@PLGA nanocomplexes showed earlier and stronger fluorescence accumulation in these organs. After 4 h post-injection, hepatic fluorescence signals gradually decreased, whereas renal signals increased, suggesting dynamic redistribution and progressive clearance of LNPmRNA@PLGA.

### Biological safety of LNPmRNA@PLGA

The biosafety of LNPmRNA@PLGA nanocomplexes was further evaluated both in vitro and in vivo. As shown in Fig. S3A, no significant cytotoxicity was observed even at an mRNA concentration of 4 μg/mL, suggesting that LNPmRNA@PLGA has good cytocompatibility. In vivo biocompatibility of LNPmRNA@PLGA was then assessed (Fig. S3B). No dose-dependent changes were detected in hematological parameters (WBC, RBC, HGB, HCT, and PLT), coagulation parameters (PT and APTT), or biochemical markers (AST, ALT, CREA, and UA). H&E staining revealed no obvious pathological alterations in major organs (heart, liver, spleen, lung, and kidney) (Fig. S3C), suggesting no apparent organ toxicity. Taken together, these findings indicate that LNPmRNA@PLGA nanocomplexes have a favorable biosafety profile.

### LIFU enhanced cellular internalization of LNPmRNA@PLGA

Because efficient intracellular delivery is essential for mRNA-based therapies for atherosclerosis, we examined whether LIFU could enhance LNPmRNA@PLGA uptake in HUVECs. HUVECs were incubated with DiI-labeled LNPmRNA@PLGA with or without LIFU irradiation, followed by confocal imaging and flow cytometry. As shown in Fig. 3A, confocal imaging revealed stronger intracellular DiI fluorescence in LIFU- treated cells than in untreated controls, accompanied by pronounced perinuclear accumulation. Consistently, flow cytometry showed that LIFU enhanced the uptake efficiency of LNPmRNA@PLGA from 78.3% to 89.3% (Fig. 3B). These findings indicate that LIFU facilitates the cellular internalization of LNPmRNA@PLGA nanocomplexes.

**Figure 3.**
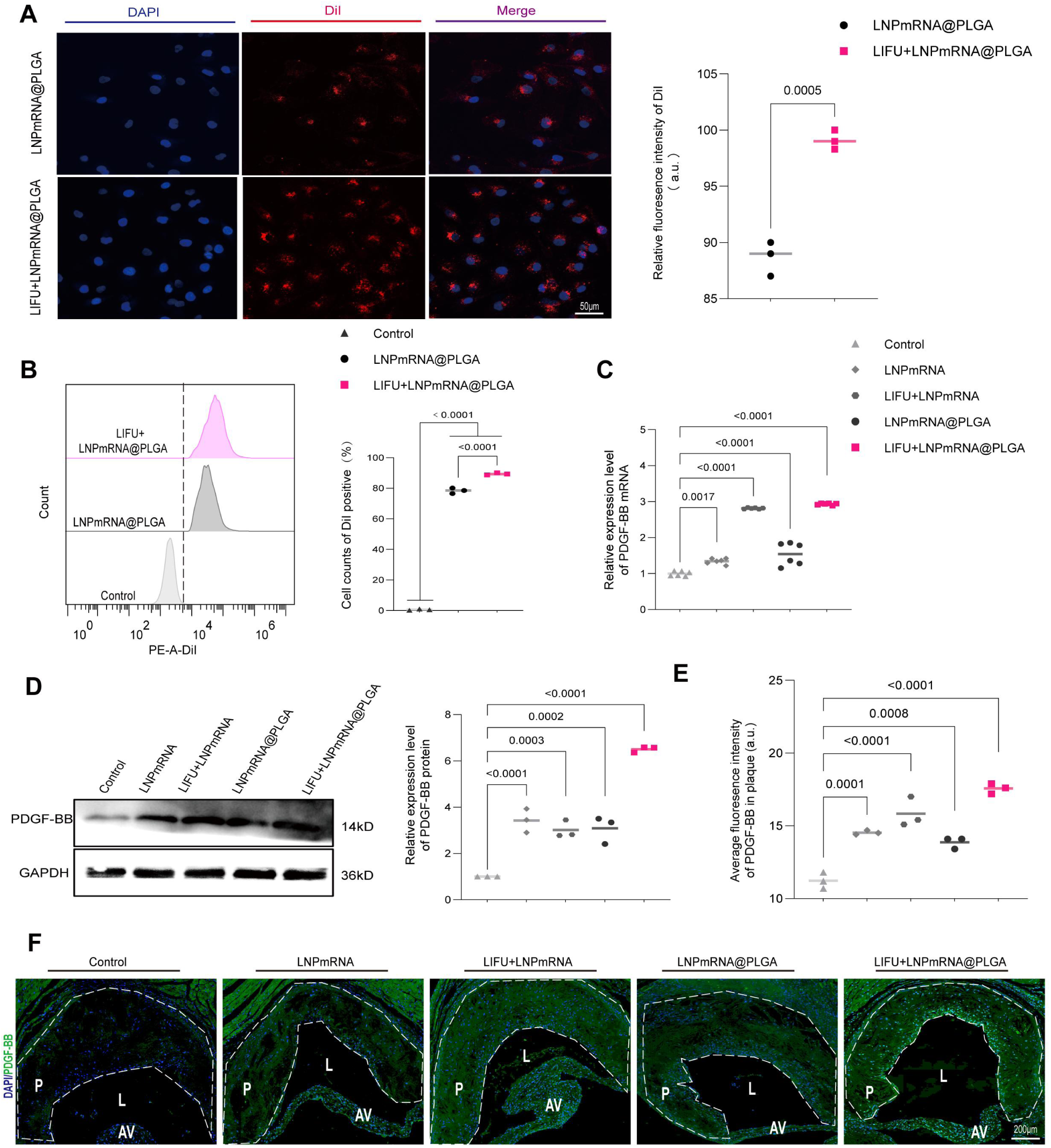
LIFU enhanced cellular uptake and PDGF-BB expression mediated by LNPmRNA@PLGA. (A) Confocal fluorescence images and (B) flow cytometry analysis of HUVECs treated with LNPmRNA@PLGA with or without LIFU irradiation. (C) PDGF-BB mRNA expression levels in HUVECs under different treatments. (D) Western blot analysis and ImageJ-based quantification of PDGF-BB protein expression in HUVECs under different treatments. (E) Statistical analysis of mean fluorescence intensity and (F) immunofluorescence images of PDGF-BB in aortic root plaques of ApoE^−/−^Fbn1^C1041G+/−^ mice. P, plaque; L, lumen; AV, aortic valve. An unpaired t test was used for (A), and one-way ANOVA was used for (B–E).

We next examined whether LIFU could enhance LNPmRNA@PLGA-mediated PDGF- BB expression in HUVECs. RT-qPCR showed slightly increased PDGF-BB mRNA expression in both LNPmRNA and LNPmRNA@PLGA groups compared with the control group. LIFU irradiation markedly amplified this effect, as evidenced by significant upregulation of PDGF-BB mRNA levels in both nanoparticle-treated groups, with the highest observed in the LIFU + LNPmRNA@PLGA group (Fig. 3C). Western blotting (Fig. 3D) confirmed a corresponding increase in PDGF-BB protein expression across the treatment groups, with the strongest signal detected in the LIFU + LNPmRNA@PLGA group (*P* < 0.0001). Consistently, IF staining further confirmed stronger PDGF-BB fluorescence in the LIFU + LNPmRNA@PLGA group (Fig. 3E, F). Overall, these data indicate that LIFU acts as a remote trigger to facilitate LNPmRNA@PLGA-mediated PDGF-BB expression.

### Paracrine effects of LNPmRNA@PLGA + LIFU treatment on vascular mural cells

Pericytes and VSMCs are major populations of vascular mural cell, and the PDGF- BB/PDGFRβ signaling pathway plays an important role in regulating their biological functions. To evaluate whether LIFU-activated nanocomplexes could modulate mural cell behavior through paracrine signaling, VSMCs and 10T1/2 cells (a pericyte precursor cell line) were cultured with conditioned media collected from HUVECs after the indicated treatments (Fig. 4A). Proliferation and migration were assessed using CCK-8 and Transwell assay, respectively. As shown in Fig. 4B, in the absence of LIFU, conditioned media from the LNPmRNA@PLGA-treated group induced weaker proliferation of both VSMCs and 10T1/2 cells than that from the LNPmRNA-treated group. After LIFU exposure, however, the proliferation rate in the two groups became comparable. Specifically, conditioned media from the LIFU-treated LNPmRNA and LNPmRNA@PLGA groups increased 10T1/2 cell proliferation to 1.9- and 1.8-fold, respectively, compared with the control group. In VSMCs, the corresponding increases were 2.6- and 2.8-fold, indicating a stronger proliferative response in VSMCs. A similar result was observed in the migration assay (Fig. 4C–E). Under LIFU irradiation, conditioned media from LNPmRNA@PLGA-treated group increased VSMC and 10T1/2 cell migration to 28-fold and 23-fold, respectively, with effects approaching those induced by media from LNPmRNA-treated group. This pattern suggests that PLGA encapsulation limits premature biological activity before LIFU activation while maintaining effective LIFU-triggered delivery, thereby promoting vascular mural cell proliferation and migration through endothelial paracrine signaling.

**Figure 4.**
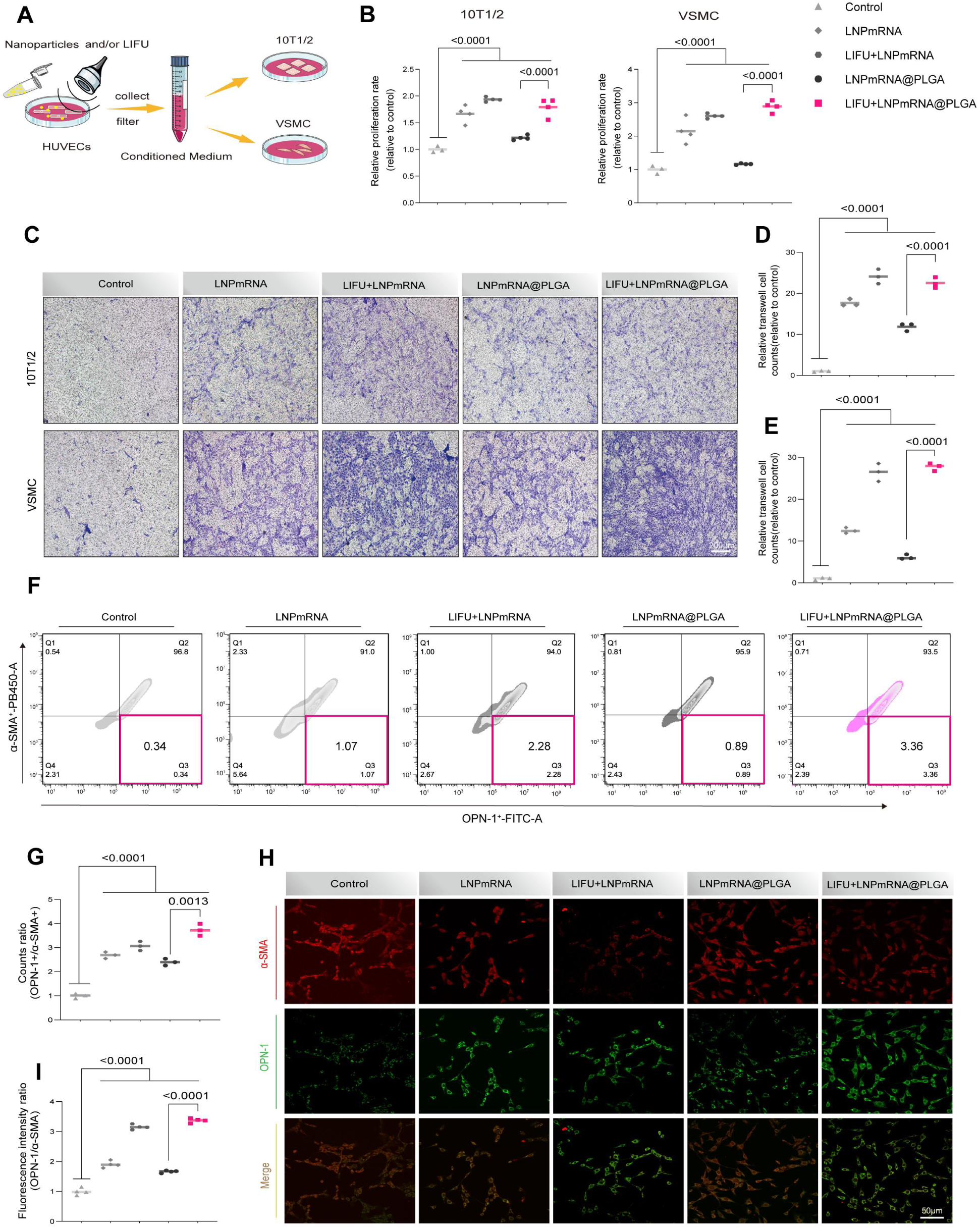
Biological effects of LIFU combined with LNPmRNA@PLGA on vascular mural cells. (A) Schematic illustration of 10T1/2 cell and VSMC treatment with conditioned media collected from HUVECs after nanoparticle treatment with or without LIFU irradiation. (B) Proliferation of 10T1/2 cells and VSMCs after treatment with different HUVEC-conditioned media. (C) Representative Transwell images and (D-E) quantitative analysis of VSMC and 10T1/2 cell migration after treatment with different HUVEC-conditioned media. (F) Flow cytometry analysis of VSMC phenotypic switching using α-SMA as a contractile marker and OPN-1 as a synthetic marker. (G) Quantification of the OPN-1+/α-SMA+ cell ratio. (H) Representative immunofluorescence images of α-SMA (red) and OPN-1 (green) expression in VSMCs. (I) Quantitative analysis of the OPN-1/α-SMA fluorescence intensity ratio. Data are presented as mean ± SEM (n = 3 independent experiments). Statistical analysis was determined by one-way ANOVA.

To further assess whether LNPmRNA@PLGA-induced endothelial paracrine signaling could mediate VSMC phenotypic switching, VSMCs were incubated with conditioned media collected from HUVECs after treatment with LNPmRNA@PLGA or LNPmRNA, with or without LIFU irradiation. α-smooth muscle actin (α-SMA) and osteopontin-1 (OPN-1) expression in VSMCs was then analyzed by flow cytometry. Flow cytometry revealed decreased α-SMA expression and increased OPN-1 expression in LNPmRNA or LNPmRNA@PLGA-treated groups. LIFU exposure further enhanced this phenotypic shift. As shown in Fig. 4F, G, after data normalization, under LIFU irradiation, the percentage of OPN-1⁺ VSMCs in the LNPmRNA@PLGA group rose to 3.7 times the level observed in the untreated control group. This phenotypic shift was further confirmed by IF staining, which showed weaker α-SMA and stronger OPN-1 fluorescence signals, particularly in the LIFU + LNPmRNA@PLGA group (Fig. 4H, I). Quantification of IF images showed that the OPN-1/α-SMA fluorescence intensity ratio reached 3.4 ± 0.3 in the LIFU + LNPmRNA@PLGA group, significantly higher than that in the control group (*P* < 0.0001). Together, these findings indicate that LIFU-triggered LNPmRNA@PLGA delivery promotes VSMC phenotypic switching toward a synthetic phenotype through PDGF-BB-associated endothelial paracrine signaling.

### Establishment and validation of a Vulnerable Plaque Mouse Model

Based on a previous report,^10^ we established a high-fat diet (HFD)-induced vulnerable atherosclerotic plaque model using ApoE^−/−^Fbn1^C1041G+/−^ mice. As illustrated in Fig. S4A, the Fbn1^C1041G+/−^ mice harbor a cysteine-to-glycine substitution at amino acid position 1041. The breeding strategy used to generate the experimental mice is shown in Fig. 5A. Fbn1^C1041G+/−^ mice were crossed with ApoE^−/−^ mice, and the offspring were further intercrossed to generate ApoE^−/−^Fbn1^C1041G+/−^ mice and ApoE^−/−^Fbn1^C1041G+/+^ mice littermate controls (Fig. S4B). All mice fed an HFD for 12 weeks before subsequent experiments.

**Figure 5.**
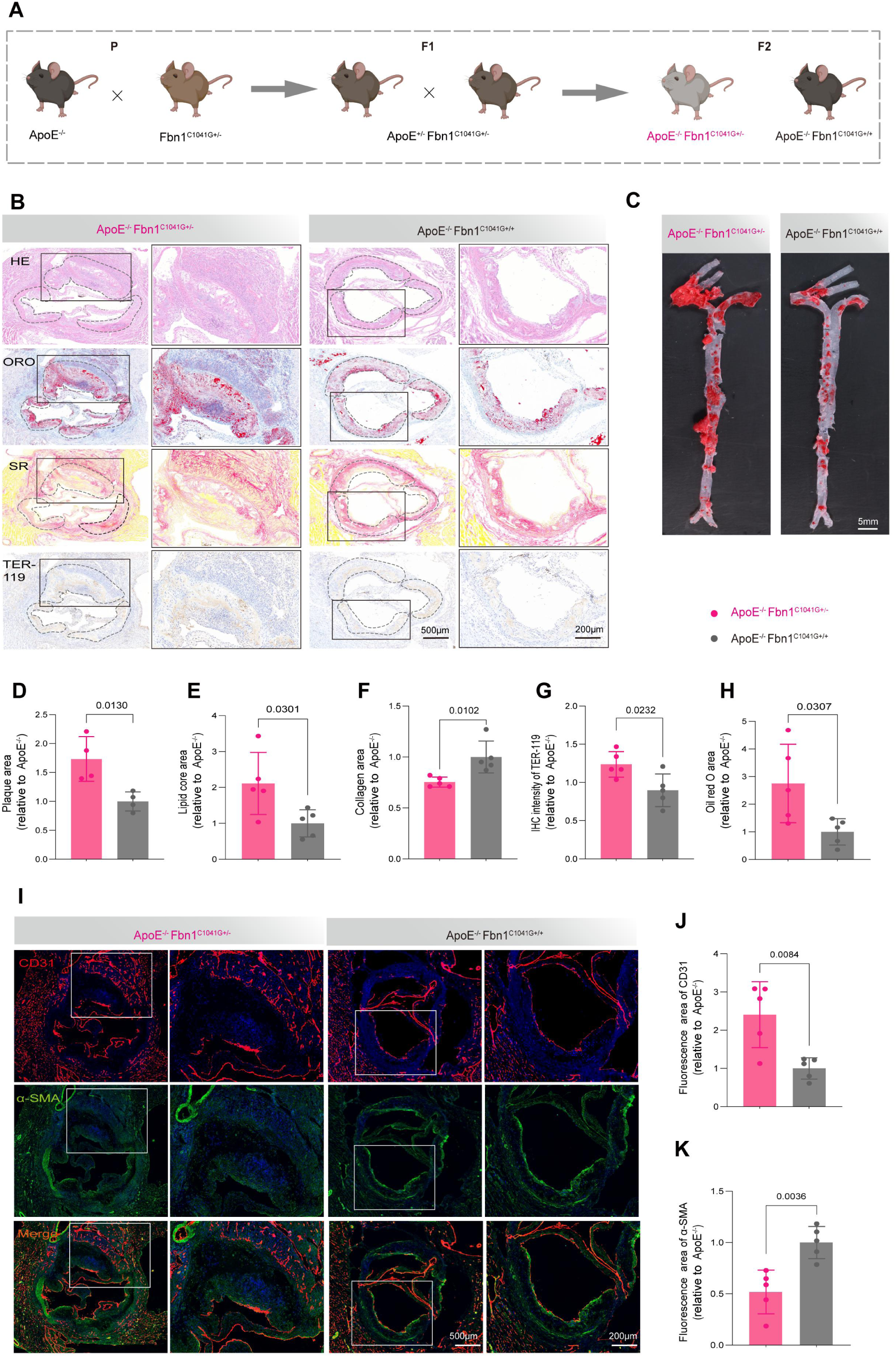
Validation of a vulnerable plaque mouse model. (A) Schematic diagram of the strategy used to construct the vulnerable plaque model. (B) H&E, Oil Red O (ORO), Sirius Red (SR), and TER-119 staining of the aortic root sections from ApoE^−/−^Fbn1^C1041G+/−^ and ApoE^−/−^Fbn1^C1041G+/+^ mice. (C) Gross ORO staining of the aorta in ApoE^−/−^Fbn1^C1041G+/−^ and ApoE^−/−^Fbn1^C1041G+/+^ mice. (D-H) Quantification of plaque area, lipid core area, collagen area, TER-119-positive area, and ORO-positive area. (I) Immunofluorescence staining for CD31 and α-SMA in aortic root sections from ApoE^−/−^Fbn1^C1041G+/−^ and ApoE^−/−^Fbn1^C1041G+/+^ mice. (J-K) Quantification of CD31+ area and α-SMA+ area. (mean ± SD, n = 5). Statistical analysis was determined by unpaired t test.

As shown in Fig. S4C, the carotid resistive index was significantly higher in ApoE^−/−^Fbn1^C1041G+/−^ mice than in the ApoE^−/−^Fbn1^C1041G+/+^ controls, suggesting altered carotid hemodynamics. Histological analysis was then performed to characterize plaque vulnerability. Aortic root plaques from ApoE^−/−^Fbn1^C1041G+/−^ mice exhibited larger lesion areas, greater lipid accumulation, reduced collagen content, and increased intraplaque erythrocyte infiltration compared with ApoE^−/−^Fbn1^C1041G+/+^ controls (Fig. 5B and D–G).

Gross ORO staining further showed a greater whole-aorta plaque burden in ApoE^−/−^Fbn1^C1041G+/−^ mice, with increased plaque number and plaque area (Fig. 5C, H). In addition, intraplaque neovascularization was frequently observed in aortic root from ApoE^−/−^Fbn1^C1041G+/−^ mice but was rarely detected in control mice (Fig. 5I-K). This was accompanied by reduced α-SMA+ VSMC coverage, indicating enhanced plaque vulnerability. Collectively, these findings indicate that ApoE^−/−^Fbn1^C1041G+/−^ mice develop plaques that recapitulate key pathological features of human vulnerable plaques, supporting their use for subsequent therapeutic studies.

### LIFU + LNPmRNA@PLGA treatment enhances plaque stability

Fig. 6A schematically illustrates the experimental timeline and therapeutic interventions. Eight-week-old ApoE^−/−^Fbn1^C1041G+/−^ mice were fed an HFD for 12 weeks to induce vulnerable plaques before treatment. The mice were then randomly assigned to three groups: the saline, LNPmRNA@PLGA, and LIFU + LNPmRNA@PLGA groups. Saline or LNPmRNA@PLGA nanocomplexes were administered intravenously once weekly for 4 weeks. In the LIFU + LNPmRNA@PLGA group, each injection was followed by LIFU irradiation at 1 W/cm² for 4 min.

**Figure 6.**
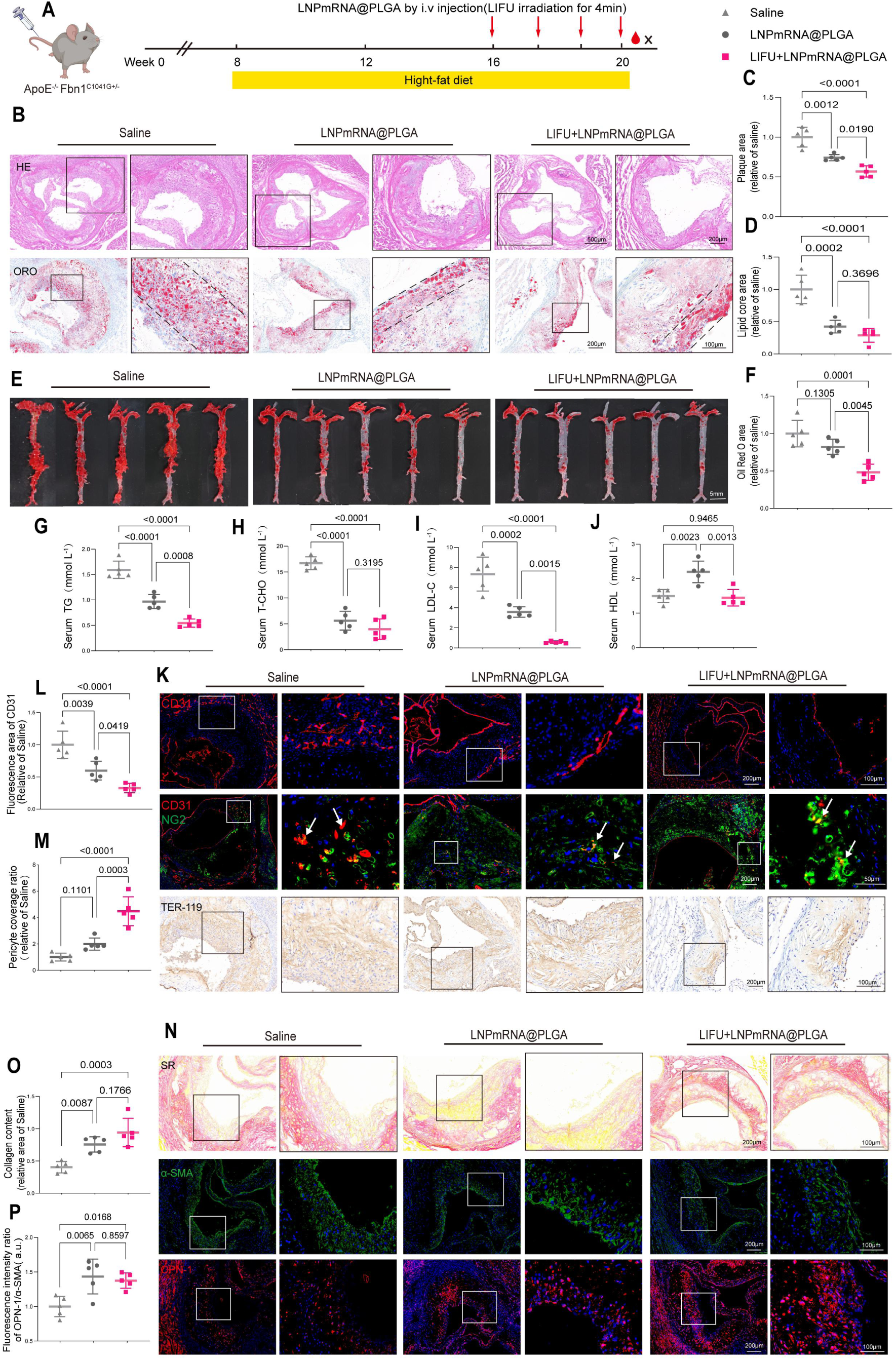
LIFU combined with LNPmRNA@PLGA enhances plaque stability. (A) Schematic diagram of the therapeutic intervention timeline in ApoE^−/−^Fbn1^C1041G+/−^ mice. (B) H&E and ORO staining of aortic root plaques after different treatments. (C-D) Quantification of plaque area and lipid core area. (E) Gross ORO staining of the aorta after different treatments. (F) Quantification of ORO-positive area. (G-J) Serum levels of TG, T-CHO, LDL-C, and HDL-C after different treatments. (K) Representative immunofluorescence staining of CD31 and NG2, and immunohistochemical staining of TER-119, in aortic root sections after different treatments. Arrows indicate intraplaque neovessels. (L-M) Quantification of the fluorescence area of CD31 and pericyte coverage ratio. (N) Sirius Red staining and Immunofluorescence staining for α-SMA and OPN-1 in aortic root sections after different treatments. (O-P) Quantification of collagen content and the OPN-1/α-SMA fluorescence intensity ratio. Data are presented as mean ± SD (n = 5). Statistical analysis was determined by one-way ANOVA.

To evaluate the plaque stabilizing effect of LNPmRNA@PLGA, atherosclerotic lesions in the aortic root were subjected to histological analysis. (Fig. 6B-D and Fig. S5). Compared with the saline group, plaque area and lipid core area were markedly reduced in the LIFU + LNPmRNA@PLGA group, with greater reductions than those observed in the LNPmRNA@PLGA group. Gross ORO staining of the aorta further confirmed a marked reduction in plaque burden in the LIFU + LNPmRNA@PLGA group (Fig. 6E, F). Serum lipid profiles were also improved after LIFU + LNPmRNA@PLGA treatment, as indicated by reduced serum triglyceride (TG), total cholesterol (T-CHO), and low-density lipoprotein cholesterol (LDL-C) levels and increased high-density lipoprotein cholesterol (HDL-C) levels (Fig. 6G-J). These findings indicate that LIFU + LNPmRNA@PLGA treatment attenuates atherosclerosis progression by reducing plaque burden, limiting intraplaque lipid deposition, and improving serum lipid profiles. We further assessed intraplaque neovessel maturation by quantifying CD31-positive neovessel area and NG2-marked pericyte coverage. As shown in Fig. 6K-M and Fig. S6A-B, CD31-positive neovessel area was significantly reduced in the LNPmRNA@PLGA and LIFU + LNPmRNA@PLGA groups compared with the saline group. In contrast, the LIFU + LNPmRNA@PLGA group exhibited markedly increased pericyte coverage, with an approximately 4-fold higher coverage ratio than the saline group, suggesting enhanced maturation of intraplaque neovessels. Consistently, reduced intraplaque hemorrhage further supported improved neovessel-associated plaque stability after LIFU + LNPmRNA@PLGA treatment (Fig. S6B). In addition, LIFU + LNPmRNA@PLGA group showed significantly increased collagen deposition and markedly increased OPN-1/α-SMA fluorescence intensity ratio in the fibrous cap compared with the other two groups, suggesting VSMC switching toward a synthetic, collagen-producing phenotype with fibrous cap reinforcement (Fig. 6N-P and Fig. S7). Overall, these findings indicate that LIFU + LNPmRNA@PLGA treatment improves plaque stability by promoting collagen deposition and VSMC phenotypic modulation, reducing excessive intraplaque neovascularization, and alleviating intraplaque hemorrhage.

### LIFU + LNPmRNA@PLGA treatment remodels the atherosclerosis microenvironment and normalizes hemodynamic alterations

To assess whether LIFU enhanced LNPmRNA@PLGA-mediated attenuation of plaque inflammation and hypoxia, CD68 and HIF-1α were analyzed as markers of macrophage infiltration and plaque hypoxia, respectively. Representative images and quantitative analysis showed that LNPmRNA@PLGA treatment reduced CD68 and HIF-1α fluorescence intensities compared with the saline group, whereas LIFU exposure further enhanced these reductions, indicating that LIFU enhanced the anti-inflammatory and anti-hypoxic effects of LNPmRNA@PLGA (Fig. 7A, B and Fig. S8A, B). Consistent with the IF results, ELISA showed that CD68 and HIF-1α protein levels in aortic plaque tissue lysates were significantly lower in both the LNPmRNA@PLGA and LIFU + LNPmRNA@PLGA groups than in the saline group, with the lowest levels observed in the LIFU + LNPmRNA@PLGA group (Fig. 7C and 7D). To determine whether these local changes were accompanied by alterations in systemic inflammation, serum TNF-α and IL-6 concentrations were measured. Both cytokines were reduced in the LNPmRNA@PLGA group and were further decreased in the LIFU + LNPmRNA@PLGA group (Fig. 7E-F). To further quantify overall plaque vulnerability, a vulnerability score was calculated as follows: (macrophage infiltration area + lipid core area)/(SMC area + collagen fiber area). Both LNPmRNA@PLGA and LIFU + LNPmRNA@PLGA treatments markedly reduced the vulnerability score, with the greatest reduction of 78.2% observed after LIFU + LNPmRNA@PLGA treatment (*P* < 0.0001, Fig. 7G).

**Figure 7.**
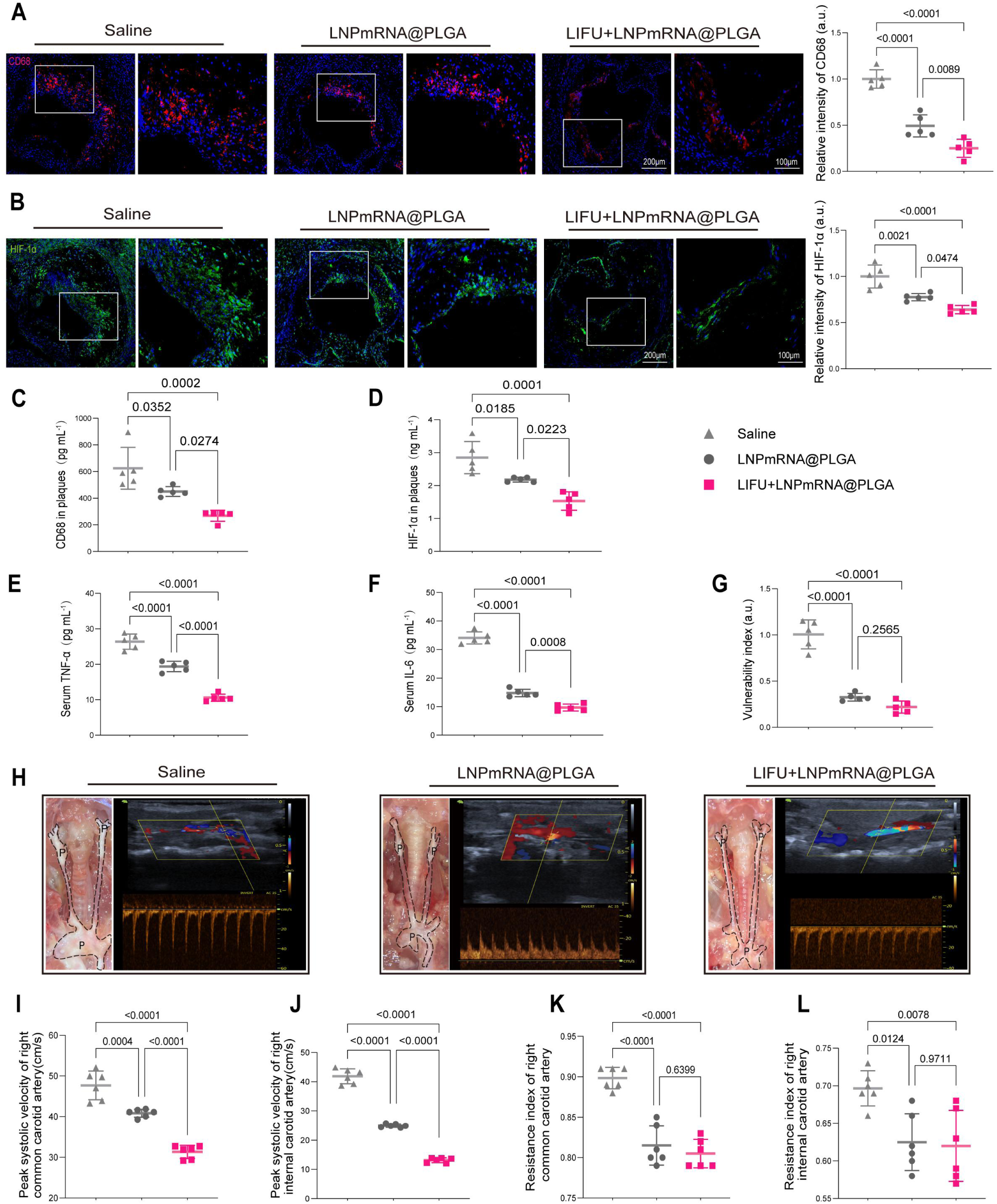
LIFU combined with LNPmRNA@PLGA improves the plaque microenvironment and carotid hemodynamics. (A-B) Immunofluorescence staining of CD68 and HIF-1α in aortic root sections from ApoE^−/−^Fbn1^C1041G+/−^ mice after different treatments. (C-D) Quantification of CD68 and HIF-1α in plaque tissues after different treatments. (E-F) Serum levels of TNF-α and IL-6 after different treatments. (G) Plaque vulnerability index in different treatment groups. (H) Color Doppler images of the right carotid artery in ApoE^−/−^Fbn1^C1041G+/−^ mice. The left panel shows the bilateral carotid arteries and corresponding plaques. P, plaque. (I-J) Peak systolic velocity (PSV) in the right common carotid artery and internal carotid artery. (K-L) Resistive index (RI) of the right common carotid artery and internal carotid artery. Data are presented as mean ± SD (n = 5). Statistical analysis was determined by one-way ANOVA.

As illustrated in Fig. 7H, plaque burden in bilateral carotid arteries and aortic arch was significantly greater in the saline group than in the LNPmRNA@PLGA and LIFU + LNPmRNA@PLGA groups. Right carotid hemodynamics were then assessed by ultrasound in each treatment group. Peak systolic velocities in the right common carotid artery and internal carotid artery were significantly reduced in the LIFU + LNPmRNA@PLGA group compared with the saline group (Fig. 7I, J), accompanied by lower carotid resistive indices (Fig. 7K, L). These results suggest that LIFU + LNPmRNA@PLGA treatment may partially normalize local carotid hemodynamics by reducing plaque burden and stenosis-associated high-velocity flow.

The integrated findings suggest that LIFU + LNPmRNA@PLGA treatment stabilizes plaques by coordinately remodeling the intraplaque microenvironment and fibrous cap structure. Reductions in excessive intraplaque neovascularization and erythrocyte accumulation suggest a lower risk of neovessel-associated hemorrhage and were accompanied by decreased macrophage infiltration and alleviated plaque hypoxia. In parallel, increased collagen deposition and VSMC phenotypic modulation toward a matrix-producing synthetic phenotype in the fibrous cap were consistent with enhanced structural stability. Together, these changes support reduced plaque vulnerability and attenuated atherosclerotic progression.

### Proteomic analysis of plaque stability following LIFU + LNPmRNA@PLGA treatment

To elucidate the molecular mechanisms underlying plaque stabilization mediated by LIFU + LNPmRNA@PLGA treatment, we performed proteomic analysis to compare the protein expression profiles between LIFU + LNPmRNA@PLGA-treated and untreated aortic tissues (Fig. 8A). A total of 367 differentially expressed proteins were identified, including 169 downregulated and 198 upregulated proteins, suggesting treatment- associated proteomic remodeling (Fig. 8B). Principal component analysis (PCA) showed that PC1 and PC2 accounted for 19% and 17.1% of the total variance, respectively, suggesting a separation trend between the treatment and control groups in the proteomic profiles of aortic tissues (Fig. 8C). Subsequent bioinformatic analyses indicated that these differentially expressed proteins were enriched in biological processes and pathways potentially associated with the stabilization of vulnerable atherosclerotic plaques. Gene Ontology (GO) enrichment analysis revealed that the differentially expressed proteins were enriched in biological processes related to platelet activation, extracellular matrix adhesion, and inflammatory responses(Fig. 8D).

**Figure 8.**
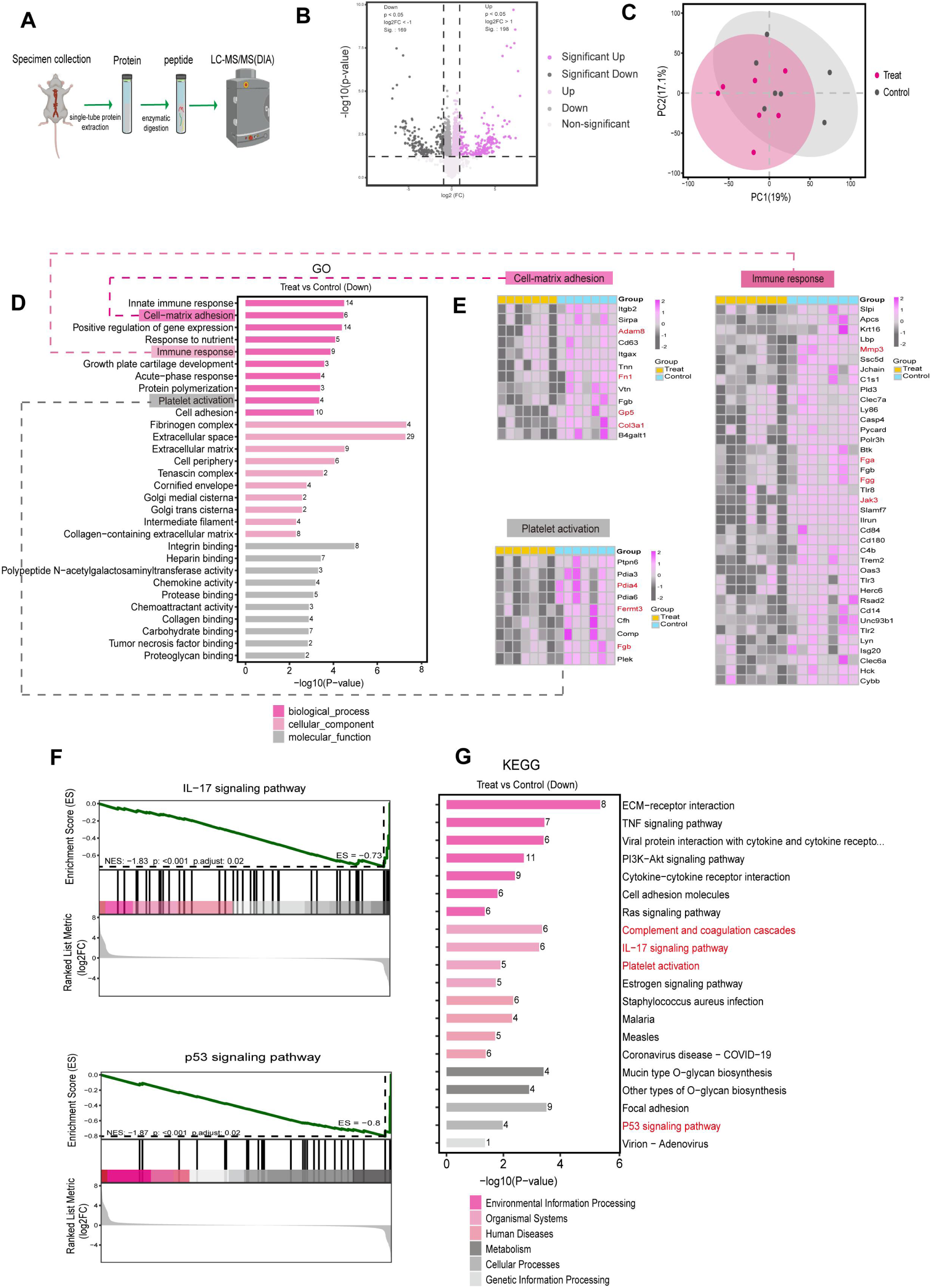
Proteomic analysis of the mechanisms underlying plaque stabilization induced by LIFU + LNPmRNA@PLGA in vivo. (A) Schematic overview of the proteomic analysis workflow. (B) Volcano plot showing differentially abundant proteins in aortic tissues from the LIFU + LNPmRNA@PLGA group compared with the control group (P < 0.05 and |log2 fold change| ≥ 1). (C) Principal component analysis (PCA) of proteomic profiles from the LIFU + LNPmRNA@PLGA and control groups. (D) Gene Ontology (GO) enrichment analysis of differentially abundant proteins between the LIFU + LNPmRNA@PLGA and control groups. (E) Heatmaps showing differentially abundant proteins associated with cell– matrix adhesion, platelet activation, and immune responses. Proteins highlighted in red represent key regulators associated with atherosclerotic plaque progression. (F) Gene Set Enrichment Analysis (GSEA) comparing the LIFU + LNPmRNA@PLGA and control groups. (G) Kyoto Encyclopedia of Genes and Genomes (KEGG) pathway enrichment analysis of downregulated proteins in the LIFU + LNPmRNA@PLGA group compared with the control group.

Heatmap visualization showed that LIFU + LNPmRNA@PLGA treatment downregulated thrombosis-related proteins, including Padia4, Fermt3, and Fgb, suggesting a potential reduction in intraplaque thrombo-inflammatory activity. In parallel, several extracellular matrix remodeling-related proteins, including Adam8, Fn1, Gp5, and Col3a1, were altered after treatment, indicating remodeling of the plaque extracellular matrix microenvironment. (Fig. 8E). In addition, downregulation of immune response-related proteins, such as Mmp3, Jak3, Fga, and Fgg was observed in the treatment group (Fig. 8E). To delineate the molecular pathways influenced by LIFU+LNPmRNA@PLGA, Kyoto Encyclopedia of Genes and Genomes (KEGG) pathway enrichment analysis was performed. 20 pathways were enriched among the downregulated proteins and were associated with the therapeutic response (Fig. 8G). Notably, the IL-17 and p53 signaling pathways, which are closely associated with atherosclerotic plaque progression, were significantly downregulated in the treatment group. Consistent negative enrichment of these pathways was further supported by Gene Set Enrichment Analysis analysis (GSEA) (Fig. 8F). These findings suggest that the therapeutic strategy may attenuate inflammatory amplification via IL-17 signaling and alleviate p53-mediated cellular stress responses, thereby contributing to a more stable plaque microenvironment. Collectively, these proteomic findings complement the histological evidence and support the plaque- stabilizing effects of LIFU + LNPmRNA@PLGA therapy.

## Discussion

Dysfunctional intraplaque neovascularization is a key driver of vulnerable plaque progression and presents a substantial challenge to current plaque-stabilizing strategies. Previous therapeutic approaches have largely focused on suppressing pathological angiogenesis; however, plaque vulnerability is determined not only by neovessel abundance but also by vessel permeability, mural cell support, and fibrous cap integrity.^1,3,10^ In this context, PDGF-BB/PDGFRβ signaling is particularly relevant because it regulates mural cell recruitment, VSMC behavior, and extracellular matrix remodeling, processes closely associated with neovessel stabilization and fibrous cap reinforcement.^10,12,13^ However, PDGF-BB has double-edged effects on vascular remodeling. Insufficient PDGF signaling may impair VSMC accumulation and fibrous cap formation, whereas excessive PDGF-BB exposure may promote undesirable vascular remodeling, intimal hyperplasia, or fibrosis.^5,10,13^ These considerations led to two key questions: first, can PDGF-BB mRNA be delivered to vulnerable plaques in a locally activatable manner? Second, can such spatially restricted expression remodel the neovessel-associated microenvironment and improve plaque stability?

In this study, we designed a LIFU-responsive LNPmRNA@PLGA nanocomplex for imaging-guided and LIFU-triggered PDGF-BB mRNA delivery to vulnerable plaques. The 7C1-based LNP component supports endothelial-oriented mRNA delivery, whereas the PFP-loaded PLGA nanobubbles provide ultrasound responsiveness and contrast- enhanced imaging capability. This design provides additional protection against premature mRNA release, while LIFU locally triggers nanocomplex dissociation, and enhances endothelial uptake and local PDGF-BB mRNA expression at the plaque site. Importantly, this platform is intended not simply to suppress intraplaque angiogenesis, but to remodel the neovessel-associated plaque microenvironment. Conditioned medium experiments suggested that LIFU-triggered PDGF-BB mRNA delivery enhanced endothelial paracrine regulation of VSMCs and pericyte cells, promoting mural cell proliferation and migration and modulating VSMCs toward a matrix-producing phenotype. In vivo, the therapeutic effects converged on three interrelated determinants of plaque vulnerability, including neovessel-associated hemorrhage, inflammatory and hypoxic stress, and fibrous cap integrity. The concurrent reduction in erythrocyte accumulation and reinforcement of the fibrous cap matrix suggest a remodeling process that shifts plaques toward a less hemorrhagic and more structurally stable phenotype, rather than a simple anti-angiogenic effect. In addition, the ultrasound imaging capability of the PLGA nanobubbles integrates ultrasound monitoring with local acoustic activation, which may be valuable for treating focally distributed atherosclerotic lesions.

Several limitations should be considered. First, although short-term biosafety was favorable, the long-term effects of repeated LNPmRNA@PLGA administration and serial PDGF-BB dosing remain to be evaluated. Defining a safe and effective therapeutic window will be essential to preserve plaque stabilization while avoiding excessive intimal growth or fibrosis. Second, although the ApoE^−/−^Fbn1^C1041G+/−^ model recapitulates several features of vulnerable plaques, it does not fully reproduce the pathophysiological features of human vulnerable plaques. Further validation in large- animal models that more closely recapitulate the pathophysiological features of human vulnerable plaques is required. Third, LIFU parameters, dosing intervals, and imaging- guided activation strategies also require further optimization. Overall, our findings provide proof of concept that ultrasound-guided, LIFU-triggered delivery enables spatially controlled PDGF-BB mRNA expression and promotes favorable remodeling of vulnerable plaques.

## Author Contributions

Q. Xie., X. Lin. and L. Deng contributed equally to this work. The majority of the experiments were conducted by them, assisted by Y. Ran., X. Yang., F. Liu., Y. Chen., J. Luo., S. Shu., D. Zhang., D. Deng., Q. Zhang., J. Ren., Z. Wang., H. Ran., and R. Huang.

Data analysis and interpretation were done by Q. Xie., X. Li., Y. Ma. and Y. Sun. The paper was prepared by Q. Xie., X. Li., L. Deng., Y. Ma. and Y. Sun.

All authors discussed the results and implications, and commented on the paper.

## Acknowledgments

The authors thank all individuals and institutional facilities that supported this work.

## Ethics Approval

All animal experiments were approved by the Ethical Committee of The Second Affiliated Hospital of Chongqing Medical University (License number: 2024–426) and conducted in accordance with the Guide for the Care and Use of Laboratory Animals. The mice were housed in a pathogen-free facility at Chongqing Medical University.

